# Repeated divergent selection on pigmentation genes in a rapid finch radiation driven by sexual selection

**DOI:** 10.1101/075713

**Authors:** Leonardo Campagna, Márcio Repenning, Luis Fabio Silveira, Carla Suertegaray Fontana, Pablo L Tubaro, Irby J Lovette

**Author notes:** To whom correspondence should be addressed. Leonardo Campagna. Cornell Laboratory of Ornithology, 159 Sapsucker Woods Road, Ithaca, NY, USA, 14850.

## Abstract

The search for molecular targets of selection is leading to a better understanding of how evolution shapes biological diversity. Instances of recent and rapid speciation are suitable for associating phenotypes with their causal genotypes, because gene flow may homogenize areas of the genome that are not under divergent selection. Locating differentiated genomic regions among taxa allows us to test associations between the genes in these regions and their contributions to phenotypic diversity. Here we study a rapid radiation of nine sympatric bird species known as southern capuchino seedeaters, which are strikingly differentiated in sexually selected characters of male plumage and song. We sequenced the genomes of 72 individuals representing a diverse set of species and associated phenotypes to search for differentiated genomic regions. We asked what genes are harbored in divergent regions and to what extent has selection on the same targets shaped phenotypic diversity across different lineages. Capuchinos show differences in a small proportion of their genomes, yet selection has acted independently on the same targets during the groups’ radiation. Many divergence peaks contain genes involved in the melanogenesis pathway, with the strongest signal originating from a regulatory region upstream of the gene coding for the Agouti-signaling protein. Across all divergence peaks, the most differentiated areas are similarly likely regulatory. Our findings are consistent with selection acting on the same genomic regions in different lineages to shape the evolution of cis-regulatory elements, which control how more conserved genes are expressed and thereby generate diversity in sexually selected traits.

## Introduction

Understanding the processes that shape biological diversity at the molecular level is a central goal of evolutionary biology. Studies of non-model organisms with divergent traits can be powerful systems in which to discover the genetic basis of distinct phenotypes. In some cases the same genes generate similar phenotypes independently in different taxa (e.g., wing color patterns in butterflies (1)). Although some phenotypic differences are caused by mutations in coding regions of causal genes (2–4), others arise through selection on areas that regulate the expression of these genes (e.g., morphological and coloration differences in African cichlids (4)). Additionally, macro-mutations such as chromosomal inversions can suppress local recombination leading to the formation of supergenes, which allow genes to co-evolve and produce complex traits (e.g., plumage coloration and mating behavior in Ruffs and White-throated Sparrows (5–7)). Despite this wealth of knowledge, connecting the evolution of phenotype to the genetic mechanisms that generate reproductive isolation and ultimately speciation remains challenging in most systems. One of the best understood cases of the genetics of speciation in animals comes from Darwin’s finches, where the morphological traits that are under selection have been identified (8), and both the molecular mechanisms that generate those traits (9–11), and the effect of trait variation on reproductive isolation are known (12, 13).

From a genomic perspective, Darwin’s finches have offered two key advantages for researchers searching for the molecular basis of the phenotypes that distinguish these birds. First, the finches have speciated recently, which translates into a relatively low background genomic differentiation. Those few areas of the genome that are highly divergent among species contain candidate loci that may have shaped the evolution of the adaptive radiation, or are still in action, even in the face of gene flow (10, 11). Secondly, there are many different species in the radiation with comparable divergence times, which leads to similarly low background genomic differentiation across multiple possible comparisons. This allows researchers to compare the genomes of more than one pair of forms with similar differences in phenotype, and assess the degree to which molecular evolution has happened in parallel (11).

The study of additional biological systems that share these tractable attributes, but which have been driven by forces other than natural selection on foraging-related phenotypes, can provide further insights into the genomics of traits that may lead to speciation. Here we focus on a group of finch-like birds from continental South America known as the southern capuchino seedeater radiation (14, 15). Capuchinos share many characteristics with Darwin’s finches (16), yet differ in that they seem to have diversified primarily via sexual selection on plumage traits that are likely melanin-based, rather than via natural selection on foraging-related traits (14, 15). Capuchino seedeaters belong to the genus *Sporophila* and are in the same family as Darwin’s finches (Thraupidae) (17), and both radiations show comparable speciation rates that are much greater than all other groups within that large family (17). The southern capuchinos are nine predominantly sympatric species that occur in Neotropical grasslands (Fig. 1A): *S. bouvreuil* (bou), *S. pileata*(pil), *S. cinnamomea*(cin), S. *ruficollis*(ruf), *S. melanogaster*(mel), *S. nigrorufa*(nig), *S. palustris*(pal), *S. hypochroma*(hypoch), and *S. hypoxantha*(hypox). Capuchinos are sexually dimorphic, and males from different species differ in secondary sexual characters such as the plumage coloration patterns they use to attract mates and the songs they use to defend their territories (14). Males defend their territories during simulated intrusions of conspecifics, but not from sympatric male capuchinos of other species (18). The species in the group are otherwise indistinguishable morphologically (19, 20) (and very similar ecologically (21, 22)), to the extent that females and juveniles lacking male secondary sexual characters cannot be identified to species even in the hand (14, 15). Despite their phenotypic diversity in male plumage, southern capuchinos show extremely low levels of genetic differentiation (14), and except for *S. bouvreuil* cannot be assigned reliably to species even using thousands of genome-wide SNP markers (15). This genetic homogeneity is a result of the groups’ recent origin during the Pleistocene, and likely the product of both incomplete lineage sorting and ongoing genetic admixture (14, 15). The apparent genetic homogeneity among southern capuchinos, despite their distinct phenotypes that are maintained in sympatry, led us to hypothesize that these species differences in male plumage may be the result of strong selection at a few key loci. We therefore sequenced the genomes of the nine southern capuchinos with the objective of locating such loci and to test whether the same targets of selection have shaped phenotypic diversity independently across different species.

**Figure 1.**
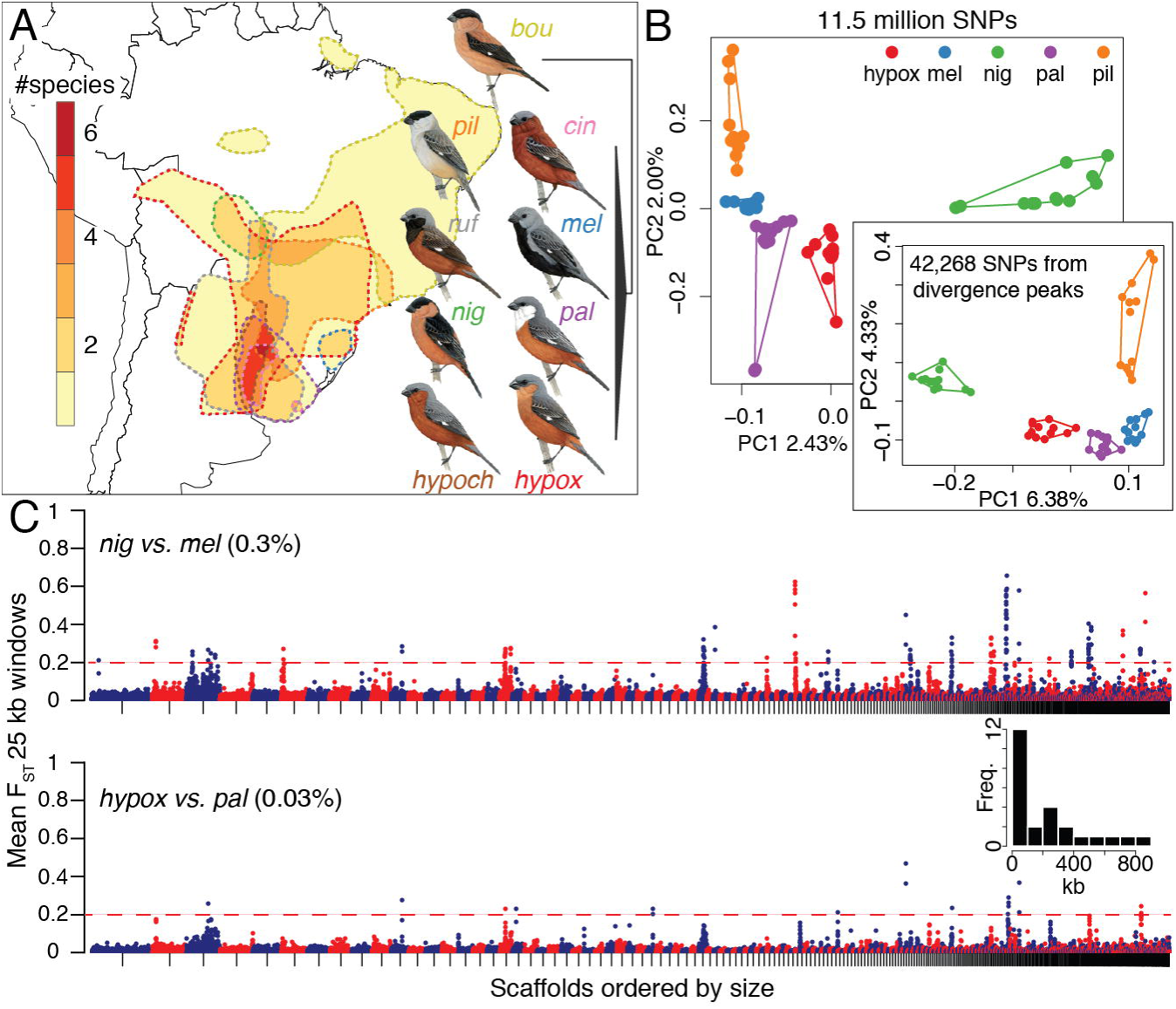
Genomic landscapes in southern capuchino seedeaters. (A) Map indicating the extent of range overlap in the nine species; note that up to six species breed sympatrically in north-eastern Argentina. The range of each species is outlined by dashed lines, with colors matching species names. The schematic phylogeny shows the relationships among species obtained from (14) and (15) (see text for name abbreviations). (B) PCA including 56 individuals of five species genotyped at ~11.5 million SNPs, and a second PCA (60 individuals) using SNPs from 442 divergence peaks alone. Four outlier individuals were omitted from the first PCA (see Fig. S1 and SI Materials and Methods for details). (C) Manhattan plots for nig vs. mel (top) and hypox vs. pal (bottom); 12 individuals per species. Each circle indicates the mean F_ST_ value for all the SNPs within a non-overlapping 25 kb window. Scaffolds in the reference genome were sorted by decreasing size and are indicated by alternating colors. The threshold for calling divergent windows is indicated by the dashed red line, and the percentage of total elevated windows is noted next to each comparison. The inset is a histogram showing the width distribution (in kb) for the 25 divergence peaks we identified.

## Results

Individual capuchinos clustered by species in a PCA derived from 11.5 million SNPs, yet the percentage of variation explained by the first two principal components was low, suggesting that a small proportion of these SNPs could be driving the pattern (Fig. 1B, Fig. S1). To search for divergent areas of the genome we compared F_ST_ values for non-overlapping 25 kb windows across the ten possible pairwise comparisons of five species (those for which our sample sizes were larger). The mean differentiation across all windows/comparisons was low (mean F_ST_ = 0.008 and SD = 0.015 across ~43.5 thousand windows), yet we found a number of divergence peaks with highly elevated F_ST_ with respect to this low background. For example, Fig. 1C shows two Manhattan plots; the upper graph (nig vs. mel) had the largest number of elevated windows (0.3%) among all comparisons and involved two allopatric species with small ranges. The bottom comparison (hypox vs. pal) had an order of magnitude fewer (0.03%) and compared two species with highly overlapping ranges. All other pairwise comparisons had a number of divergence peaks that ranged between the extremes shown in Fig. 1C (Fig. S2). We identified a total of 25 divergence peaks with elevated F_ST_ (>0.2) that are candidate targets of selection driving species differences among capuchinos (Table 1); these peaks range in width from 25 to 840 kb (average ∽243 kb, inset in Fig. 1C). SNPs from these divergence peaks alone can be used to assign individuals to species in a PCA (inset in Fig. 1B). We found 99 SNPs that were fixed (F_ST_=1) in at least one comparison across all pairwise combinations of five capuchinos; these represented 65 different sites in the genome. Because our sample sizes were low for the remaining capuchino species (bou, cin, hypoch, ruf), we did not use them to identify divergence peaks. A PCA using the SNPs from under the peaks showed some overlap between taxa when we included all nine species (Fig. S1, mainly between ruf vs. hypox and bou vs. pil). It is therefore possible that there are some additional undetected areas of the genome that are involved in capuchino seedeater differentiation.

Next we asked if the same divergence peaks were involved in differentiation across multiple combinations of capuchino species in ways that imply independent patterns of selection on the same loci. The left panels in Fig. 2 show examples of divergence peaks (5 kb windows) with the ten different pairwise comparisons overlaid. Although no single area of the genome (i.e., differentiation peak) was present in all comparisons, many are found in multiple comparisons. For example, the peak in Fig. 2A was present in nine out of the ten comparisons, many of which involved independent pairs of species (e.g., nig vs. mel and pil vs. hypox). Other divergence peaks were less ubiquitous, yet are present across multiple pairs of species (left panels in Fig. 2, Fig. S3, and Fig. S4). To better understand the nature of the differences among species within the divergence peaks, we conducted PCAs with the SNPs from each of these areas separately. Fig. 1B shows four clusters of haplotypes in the region under the most common peak in our dataset. Other divergence peaks varied in the species they were present and the extent to which they could be used to diagnose species in PCAs (center panels in Fig. 2, Fig. S3, and Fig. S4).

**Figure 2.**
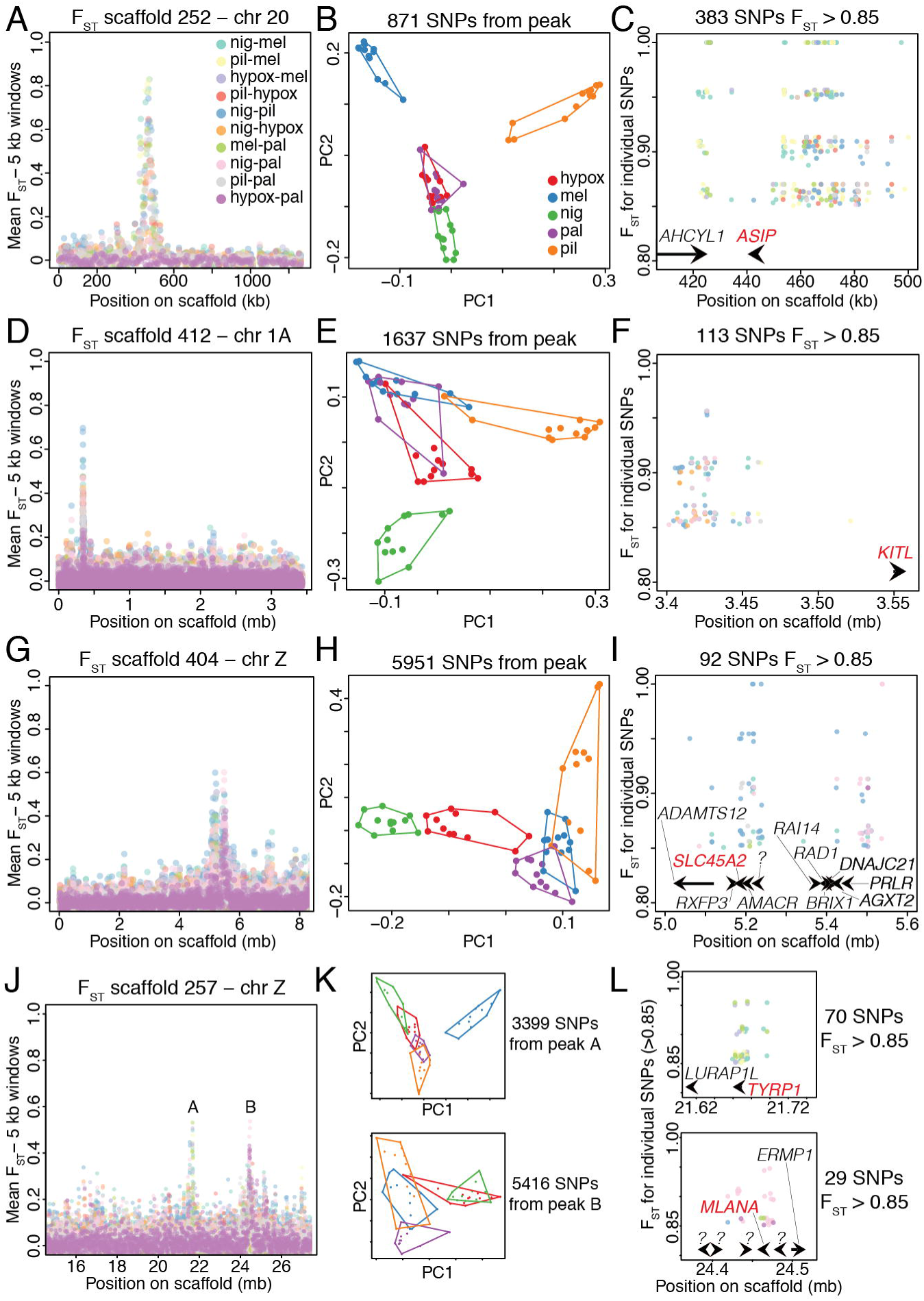
Repeated selection on pigmentation genes in different capuchino species. (A) The divergence peak on scaffold 252, which mapped to chromosome 20 in the Zebra Finch. The ten possible pairwise comparisons across five capuchino species are overlaid (see color-code legend to identify specific comparisons). Each circle is the mean F_ST_ value for all SNPs within a non-overlapping 5 kb window. (B) PCA for 60 individuals of 5 species using the SNPs from under the peak in (A); see the color-code in the legend to identify the species. (C) F_ST_ and genomic location of individual SNPs with values of 0.85 and higher, color-coded by pairwise comparison as in (A). The positions of genes that are close to these highly divergent SNPs are indicated by arrows drawn to scale. Names in red note genes involved in the melanogenesis pathway. (D, E, and F) As above, for the divergence peak on scaffold 412. (G, H, and I) As above, for the divergence peak on scaffold 404. (J, K, and L) As above, for the divergence peak on scaffold 257. (K and L) The top plot corresponds to the peak labeled “A” and the bottom one to the peak labeled “B” in (J). Annotations with question marks did not match genes with known names.

We identified a total of 246 gene models within these divergent areas of the genome (an average of 10 per divergence peak), 156 of which matched genes of known functions in other species. We performed an enrichment analyses to understand if genes in this list were predominantly involved in certain pathways. The most prominent hit was the melanogenesis pathway (KEGG pathway analysis, p= 2.0E-3). We found nine melanogenesis genes in eight different divergence peaks (Table 1). The peak containing the gene coding for the Agouti-signaling protein (ASIP) had the largest number of highly divergent SNPs in our dataset: 30% of all observed SNPs with F_ST_ >0.85 and 58% of all SNPs with F_ST_=1 (Fig. 2C). Accordingly, this was the peak that showed the greatest increase in absolute sequence divergence (measured using the Dxy statistic) when comparing the region under the peak to the areas on the same scaffold outside of the peak (Fig. S5). Other peaks contained a smaller number of these highly divergent SNPs (Fig. 2, Table 1, Fig. S3, Fig. S4). The peaks containing melanogenesis genes accumulated 60% of SNPs with F_ST_>0.85 and 63% of fixed SNPs (across all pairwise comparisons). Fig. 2 and Fig. S3 show the eight divergence peaks containing melanogenesis genes, ranked from top to bottom by the number of highly divergent SNPs (F_ST_>0.85) present in these peaks (summarized in Table 1). The right panels in these figures indicate the position of these SNPs with respect to the closest gene models. We also identified three peaks that together accumulated 30% of the observed highly divergent SNPs and did not contain melanogenesis genes (Fig. S4), however one of these peaks contained the gene HERC2. An intron within this gene functions as an enhancer which regulates the expression of the OCA2 pigmentation gene in humans, involved in controlling eye, hair and skin color (23). The remaining peaks in Fig. S4 did not contain genes that could be easily associated to plumage or other phenotypes (Fig. S4). The remaining 14 peaks (Table 1) accounted for only 10% of the observed highly divergent SNPs. Finally, because the Melanocortin 1 Receptor (MC1R) is known to affect plumage coloration in many bird species and interact directly with ASIP (24), we asked if this gene showed divergence among capuchino seedeaters and had been overlooked in our analysis. MC1R was present in our reference genome assembly, yet did not show differences in any of the pairwise comparison across our five capuchino species (Fig. S6).

Nearly all the fixed sites we observed (99%) were located in non-coding areas of the genome. The most common peak in our dataset concentrated 58% of these fixed differences within several thousand kb up and downstream of the ASIP gene (Fig. 2C). We found that these areas contained positions that were highly conserved across the genomes of distantly related birds (Turkey, Chicken, Budgerigar and Zebra finch), comparable in their levels of conservation to certain positions on the exons of ASIP (Fig. S7). It is therefore likely that these regions contain cis-regulatory elements that control the expression of ASIP, and a similar situation may be true for the other differentiated regions found in close proximity to genes.

## Discussion

Despite their striking differences in male plumage, southern capuchino seedeaters are differentiated only in a small proportion of their genomes. The identity of these rare differentiated genomic regions differs somewhat among capuchinos, yet in many cases the same divergent regions are present in comparisons across many pairs of species. This convergence of differentiation across multiple independent pairs of species implies that selection has acted repeatedly in different lineages on the same genomic targets to shape phenotypes. Many of the most highly differentiated areas of the capuchino genome contain genes that are part of the melanogenesis pathway. The area upstream of the gene coding for the Agouti-signaling protein is the most ubiquitous peak, showing the strongest signal of differentiation. More generally, we observe narrow divergence peaks in different components of the melanogenesis pathway that are genetically unlinked. Plumage coloration is generally important for reproductive isolation in birds (25), and differences in genes that control melanin-based variation in plumage have been found in different pairs of incipient avian taxa (e.g., carrion and hooded crows (26), flycatchers of the Solomon Islands (27), blue-winged and golden-winged warblers (28)). The differences we observe in capuchinos are mostly in non-coding areas of the genome, therefore our findings are consistent with the evolution of cis-regulatory elements. These regulatory elements may vary by species, yet control the expression of the same set of genes, generating the strikingly different phenotypes we observe in the capuchinos. In particular, the regulation of the expression of melanogenesis genes may also lead to pigmentation differences across the plumage patches within each species (e.g., throat vs. back) (Fig. 1A).

Three factors could have contributed to the rapid evolution of phenotypic diversity in the capuchinos. First, a very large effective population size was inferred for the ancestor of the radiation (15). The amount of genetic variation a population can sustain is proportional to its size, with large populations providing more possible substrates for rapid evolution from standing genetic variation to take place (29). Additionally, differences could have accumulated among species in allopatry and eventually been exchanged via hybridization, leading to novel phenotypes as has been described for *Heliconius* butterflies (30). In particular, the differentiation and exchange via hybridization of regulatory elements that control the expression of more conserved genes has been found to drive phenotypic diversity in *Heliconius*(31). A similar situation could have contributed to phenotypic diversity in the capuchinos, which also show modular variation in their plumage, with the same patches having different colors depending on the species (e.g., throat, back or belly can be black, white, cinnamon or rufous in different species; Fig. 1A). Finally, 10 out of the 25 divergence peaks that we identified are located on the Z chromosome. Sex chromosomes are known to evolve faster than autosomes because of their smaller effective population size (32). Moreover, sex chromosomes may also be particularly relevant to speciation (33), as suggested by Haldane’s rule, where the heterogametic sex is commonly inviable or missing from hybrid crosses (34). For both reasons divergence in genes located on sex chromosomes could have facilitated rapid evolution in capuchinos.

We have identified genomic substrates that lead to phenotypic diversity in capuchinos, differences that are likely relevant to mate recognition and eventually reproductive isolation in these strongly sexually dimorphic species. Because the most divergent areas across the genomes of capuchinos contain pigmentation genes, this leads to the question of whether we have found the genes responsible for maintaining these lineages as separate species. The pigmentation genes are linked to other loci for which the connection between genotype and phenotype is harder to make, and we do not understand the contribution of these additional genes to species differences. It is possible that prezygotic isolation via mate choice is strong enough to maintain the phenotypic integrity of capuchino species even though many species breed in local-scale sympatry. However, we also cannot at this point discard the possibility that postzygotic incompatibilities exist between species, perhaps associated with these same divergent regions of the genome. As natural selection has shaped the beaks of finches in the Galapagos, leading to the generation of biological diversity, our study suggests sexual selection may have shaped the plumage and songs of male capuchinos, generating yet another extraordinary rapid radiation of finch-like birds.

## Materials and Methods

### Reference genome assembly and annotation

We combined short-read Illumina data (estimated depth of coverage of 112x) with long-read Pacific Biosciences sequencing (estimated depth of coverage of 9.1x) to assemble the genome of a *S. hypoxantha* individual using ALLPATHS-LG (35) and PBJelly (36). The total length of the assembly was 1.17 Gb distributed in 5120 scaffolds, with an N50 of 8.7 Mb and 4.8% Ns. Our reference genome contained a single copy of 90% of a set of conserved vertebrate genes used to assess assembly quality (37). We annotated the *S. hypoxantha* genome with MAKER (38), which produced a total of 14,667 gene models (75.5% of the 19,437 genes annotated in the Zebra Finch genome). We searched for annotations of interest in the UniProt database (http://www.uniprot.org/) and identified enriched pathways using DAVID v6.7 (39). See SI Materials and Methods for details.

### Population level sequencing and variant discovery

We re-sequenced the genomes of 72 individuals from nine capuchino species. We included 12 individuals per species for nig, pil, mel, hypox, and pal, and 3 individuals per species for bou, cin, hypoch, and ruf. After quality filtering we aligned individually barcoded samples to the reference genome using Bowtie2 (40). Because *S. hypoxantha* shows similar genetic distances to all other members of the clade (14),we did not observe a species effect on mapping efficiency or quality. We called SNPs following the GATK best practices (41). After filtering low quality variants we retained 11,530,110 SNPs (referred to as 11.5 M SNPs), genotyped for 72 individuals of the nine southern capuchino species (mean depth of coverage per species: nig 3.8x, pil 4.1x, mel 4.8x, hypox 4.3x, pal 5.7x, bou 7.0x, cin 4.1x, hypoch 5.9x, ruf 5.2x). See SI Materials and Methods for details.

### Population genomic analyses

We calculated F_ST_ values for non-overlapping 25 kb and 5 kb windows, and for individual SNPs using VCFtools (42). These statistics were calculated for the ten different pairwise comparisons across the five capuchinos with larger sample sizes (nig, pil, mel, hypox, and pal). We identified divergence peaks using the average F_ST_ value for 25 kb windows, discarding regions with less than two windows and windows with less than 10 SNPs. The species with smaller sample sizes were not used to identify divergence peaks. We selected regions that showed an F_ST_ value elevated above 0.2 (between 12 and 13 standard deviations above the overall F_ST_ mean). We only retained candidate regions that had at least one individual SNP with an F_ST_ of 0.85 or higher, which we considered a putative target of selection. In total, we identified 25 divergent regions using these criteria. See SI Materials and Methods for details

## Acknowledgements

This project was funded by NSF grant DEB – 1555754 to IJL, by PICT 2014-2154 (ANPCyT, Argentina) to PLT and by a Genomics Scholar fellowship from the Center for Vertebrate Genomics (Cornell University) to LC. We thank the members of the Fuller Evolutionary Biology lab group (particularly B Butcher, D Toews, S Taylor, N Mason, S Aguillon, J Walsh-Emond, P Dean-Coe, N Hofmeister, J Berv and B Van Doren), C Dardia, J Lewis, D Lijtmaer, and R Razavi for support and feedback on this project. We thank government agencies of Brazil (CNPq, CAPES-PROEX, FAPESP, ICMBio, SISBIO 36881-1, and CEMAVE 361788) and Argentina for grants and permits. For tissue loans, we are indebted to the Museo Argentino de Ciencias Naturales “Bernardino Rivadavia” (MACN, CONICET, Argentina), University of Kansas Natural History Museum (USA) and Museu de Ciências e Tecnologia of the Pontifícia Universidade Católica do Rio Grande do Sul (Brazil). Illustrations in Fig. 1A were obtained with permission from (43).

